# Image Processing in the Acute to Chronic Pain Signatures (A2CPS) Project

**DOI:** 10.1101/2024.12.19.627509

**Authors:** Patrick Sadil, Konstantinos Arfanakis, Enamul Hoque Bhuiyan, Brian Caffo, Vince D Calhoun, Daniel J Clauw, Mark C DeLano, James C Ford, Ramtalik Gattu, Xiaodong Guo, Richard E Harris, Eric Ichesco, Micah A Johnson, Heejung Jung, Ari B Kahn, Chelsea M Kaplan, Nondas Leloudas, Martin A Lindquist, Qingfei Luo, Todd A Mulderink, Scott J Peltier, Pottumarthi V Prasad, Christopher Sica, Joshua Urrutia, Carol GT Vance, Tor D Wager, Yang Xuan, Xiaohong J Zhou, Yong Zhou, David C Shu, The Acute to Chronic Pain Signatures Consortium

## Abstract

The Acute to Chronic Pain Signatures (A2CPS) project is a large-scale, multi-site initiative aimed at identifying biomarkers and biosignatures that predict the transition from acute to chronic pain. The project is collecting multimodal, longitudinal data from over 2,500 individuals at risk for developing chronic pain after surgery. Here we describe the neuroimaging component of A2CPS, including the acquisition protocols, processing pipelines, and contents of the initial data release. The imaging protocol includes structural, diffusion, resting-state and task-based functional magnetic resonance imaging (MRI) data. Data are collected across multiple clinical sites using different scanner manufacturers, with attention to protocol harmonization and quality control. The processing pipeline integrates several established neuroimaging tools to extract potential biomarkers, including measures of brain structure, connectivity, and pain-related neural signatures. The first data release includes pre-surgical imaging data for 595 participants, with high quality ratings across modalities (98.7% of sMRI, 99.8% of dMRI, and 94.6% of fMRI images were rated as acceptable or better). Initial analyses demonstrate expected relationships between brain-derived measures and clinical variables, such as associations between brain age and psychological factors. This dataset represents a valuable resource for both pain research and neuroimaging methods development, with future releases planned to include additional participants and expanded analysis pipelines and processed data derivatives.

## I. Introduction

For most people, pain due to tissue damage or inflammation decreases as the injury heals. However, sometimes the pain of an injury, surgery, or disease can linger and become chronic. Within the United States, chronic pain has been a contributing factor to the current opioid epidemic^1^, and it is also recognized as a distinct, widespread public health issue affecting tens of millions of people directly^2–4^. The causes, risk factors, and mechanisms of pain chronification remain unclear^3–5^.

To catalyze discoveries in pain chronification, the US National Institute of Health Common Fund initiated the Acute to Chronic Pain Signatures (A2CPS) project^6,7^, a multi-disciplinary endeavor to develop biomarkers and biosignatures of pain chronification after knee or thoracic surgery. The project aims to test candidate, prospective biomarkers (univariate factors that prior evidence indicates may be related to chronic pain transition)^8–10^, assemble putative biomarkers into biosignatures (measures derived from combinations of several biomarkers), and catalyze discovery of novel biomarkers or biosignatures. Many factors have been found to be relevant to the transition to chronic pain, including psychosocial, brain, genetics, genomics, and metabolic pathways ^11–13^, so A2CPS casts a wide net across multiple modalities (Figure 1; Table 1). In this report, we describe the neuroimaging data collected in the project, including the acquisition, preprocessing, and quality control procedures in the first public data release of A2CPS (Release 1.1.0).

**Figure 1.**
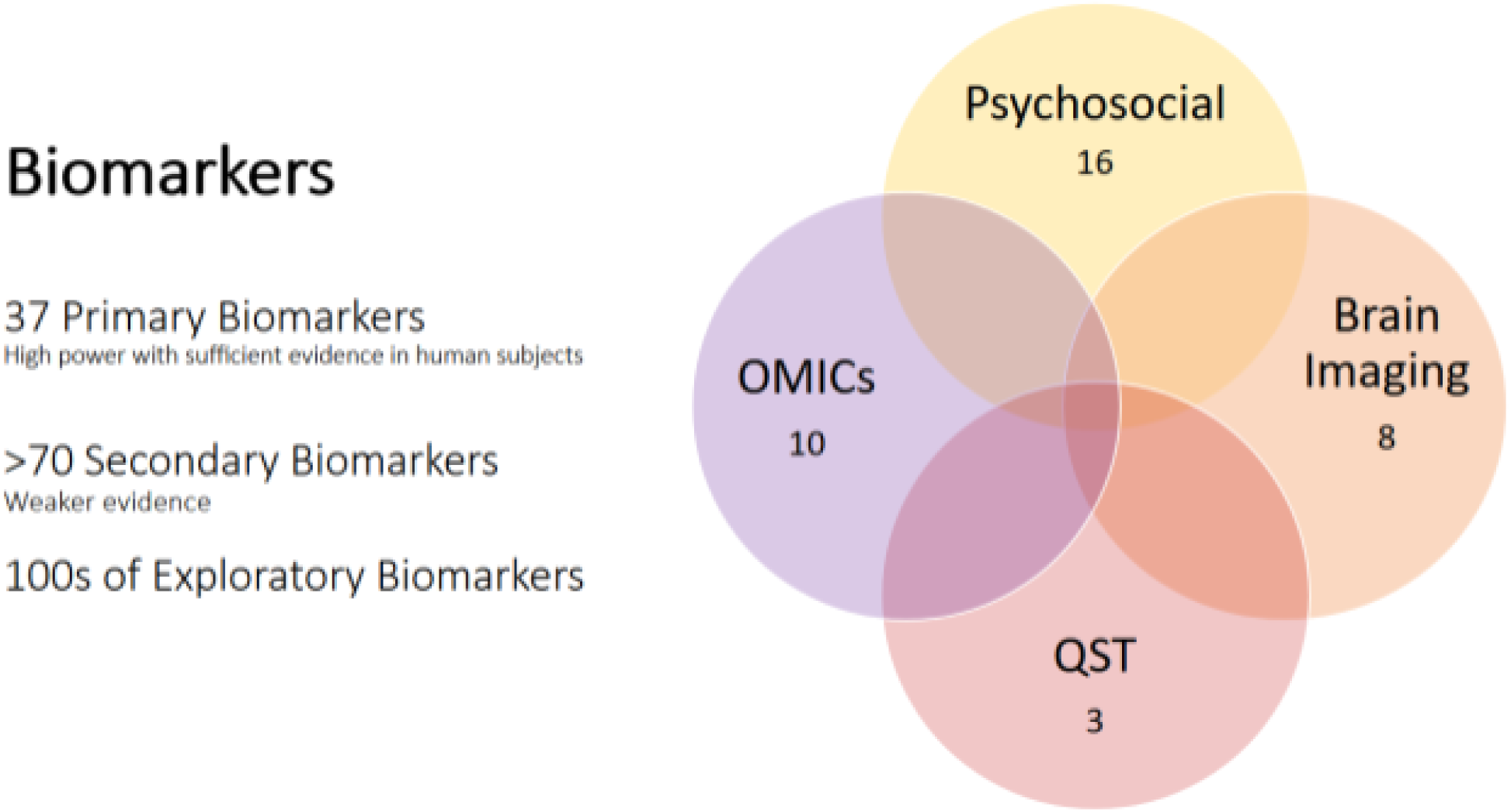
A2CPS biomarkers provide overlapping coverage of chronic pain effects and mechanisms.

**Table 1.** Primary and Secondary A2CPS Imaging Biomarkers.

| <u>Primary Biomarkers</u> | <u>Secondary Biomarkers</u> |
| --- | --- |
| sMRI – Volume of medial prefrontal cortex<br>dMRI – Structural integrity in nucleus accumbens, medial prefrontal cortex<br>rsfMRI – Core DMN; vmPFC-NAc functional connectivity<br>rsfMRI – Core DMN; DMN network–S1 functional connectivity<br>rsfMRI – Core DMN; DMN-Insula Connectivity<br>rsfMRI – Hub disruption in target network<br>tfMRI – Evoked Response in Neurologic Pain Signature (NPS)<br>tfMRI – Evoked response in fronto-striatal systems related to descending/central pain modulation and self-regulation (SIIPS) | sMRI – T1 gray matter density<br>sMRI – T1 gray matter morphometry<br>sMRI – Volume of hippocampus<br>sMRI – Volume of amygdala<br>rsfMRI – Node degree in target regions<br>rsfMRI – Lateral thalamus functional connectivity with periaqueductal gray, brain functional marker for CPM<br>dMRI – White Matter Tractography – FA in superior longitudinal fasciculus (fronto-parietal), internal and external capsule (corticostriatal)<br>tfMRI – Local pattern responses in nociceptive regions<br>tfMRI – Local pattern responses in context-related regions |

Neuroimaging, and in particular magnetic resonance imaging (MRI), has provided a rich medium for developing potential biomarkers for chronic pain^14–17^. A full review is beyond the scope of this article, but here we highlight the major kinds of measures that may be usefully derived from imaging. Incidences of chronic visceral pain, musculoskeletal pain, and migraine have been characterized by neuroanatomical structure^18,19^ and structural connectivity^20^. Many studies have identified patterns of functional connectivity that differentiate cases (individuals with or who develop chronic pain) and controls^5,21–25^, as have higher-order graph theoretic properties^26–28^. Current and future pain is also related to functional MRI (fMRI) activity during the experience of pain and other emotionally and motivationally relevant stimuli^29–31^. Beyond regional activation, individuals with chronic pain can also be classified from controls based on whole-brain patterns of neural activity in response to tonic pain stimuli^32,33^. As a whole, the literature suggests structural and functional information, both at rest and in response to painful stimuli, may contribute informative biomarkers for chronic pain.

To have sufficient statistical power to test an expansive list of biomarkers requires a large dataset, and so the A2CPS project will include over 2500 participants. Data will be released in stages, with the first public release available by Winter 2025 on the US National Institute of Mental Health Data Archive (NDA). The participants are undergoing either total knee replacement or thoracic surgery, which serves as a tightly-controlled acute injury. Although many participants are expected to recover from surgery, for a subset, the pain of that injury will likely become chronic^34–37^. Participants are scanned before their surgery, and a subset will be scanned again 3 months after the surgery, when pain chronification can be diagnosed^38^. In consideration of the existing candidates for imaging biomarkers, the imaging protocol includes structural MRI (sMRI), diffusion MRI (dMRI), resting-state functional MRI (rfMRI), and task functional MRI (tfMRI). For the neuroimaging research community, the data and associated pipelines open a new avenue for studying predictive biomarkers, and to help address the pressing health issue of chronic pain.

## II. Data Acquisition

Imaging-related activities in the A2CPS project are divided among two multisite clinical centers (MCCs) where image acquisitions occur and a Data Integration and Resource Center (DIRC) which oversees image processing and analysis^10^. The A2CPS consortium also includes a Clinical Coordinating Center (CCC), three omics centers, and representatives from the NIH Pain Consortium, Common Fund, and National Institute of Drug Abuse.

### Acquisition overview

The A2CPS MRI protocol is based on the imaging protocol used in the Adolescent Brain Cognitive Development (ABCD) study^39^, but adapted to consider the unique constraints involved in scanning perioperative patients and with modifications tailored to the design and interests of the study and to the scanner hardware of participating collection sites. The selection of sites for the A2CPS project was constrained by the ability to access appropriate patient populations in sufficient numbers. This necessitated a multi-site image collection approach, and furthermore one that would include MRI scanners from multiple manufacturers. The two A2CPS Multisite Clinical Centers, MCC1 and MCC2, use a mix of General Electric, Siemens, and Philips scanners (Table 2), all of which were supported under the ABCD imaging protocol. Note that MCC2 entered the project a year after MCC1, and thus as of Release 1 the majority of image acquisitions are from MCC1.

**Table 2.** Sites, site codes, and information on scanners, multiband sequences, and head coils. Note that some site names have changed since codes were set, and that the MCC2/UM site uses two scanners. Scanner software versions were as listed for all scans of Release 1, but updates to software and hardware are expected for subsequent releases. RU joined the consortium after the first data freeze and is contributing scans to subsequent releases.

| **Multisite**<br>**Clinical**<br>**Center** | **Site** | **Vendor** | **Model &**<br>**Scanner**<br>**Bore (cm)** | **Version** | **Head Coil**<br>**Channels** | **Multi-band** | **Scans**<br>**(R1)** |
| --- | --- | --- | --- | --- | --- | --- | --- |
| MCC1 | U. Illinois<br>Chicago<br>(UI) | GE | Discovery<br>MR750,<br>60 | DV26.0 R02<br>1810.b | 32 | ABCD/GE | 179 |
|  | Endeavor<br>Health<br>(NS) | Siemens | Skyra Fit,<br>70 | syngo MR<br>E11 (VE11C) | 64 | MGH | 169 |
|  | U.<br>Chicago<br>(UC) | Philips | Ingenia<br>dStream,<br>70 | 5.6.1.0 | 32 | Philips | 68 |
| MCC2 | U.<br>Michigan<br>(UM) | GE | Discovery<br>MR750,<br>60 | DV26.0 R02<br>1810.b | 32 | ABCD/GE | 134 |
|  |  | GE | Signa<br>UHP, 60 | RX28.0 R04<br>UHP3T<br>2111.a | 32 | ABCD/GE |  |
|  | Wayne<br>State<br>(WS) | Siemens | Verio, 70 | syngo MR<br>B17 | 32 | CMRR | 22 |
|  | Corewell<br>Health/MS<br>U - Grand<br>Rapids<br>(SH) | Siemens | Magneto<br>m Prisma,<br>60 | syngo MR<br>E11 (VE11C) | 64 | Siemens | 23 |
|  | Rush<br>University<br>(RU) | Siemens | Magneto<br>m Prisma,<br>60 | syngo MR<br>XA30 | 64 | Siemens | 0 |

We have created a data rich yet time-efficient (approximately 45 minutes) MRI protocol comprising (Figure 2): (a) structural MRI (sMRI) with T1-weighted contrast; (b) diffusion-weighted imaging (dMRI) with multi-shell, multi-directional acquisitions; and (c) functional MRI (fMRI) before, during, and after a tonic painful stimulus. To enable correction of magnetic susceptibility distortion artifacts in both the diffusion and functional images, the protocol also includes pairs of susceptibility weighted images (swMRI): spin echo echo-planar imaging scans with opposite phase encoding directions matching the dMRI and fMRI images.

**Figure 2.**
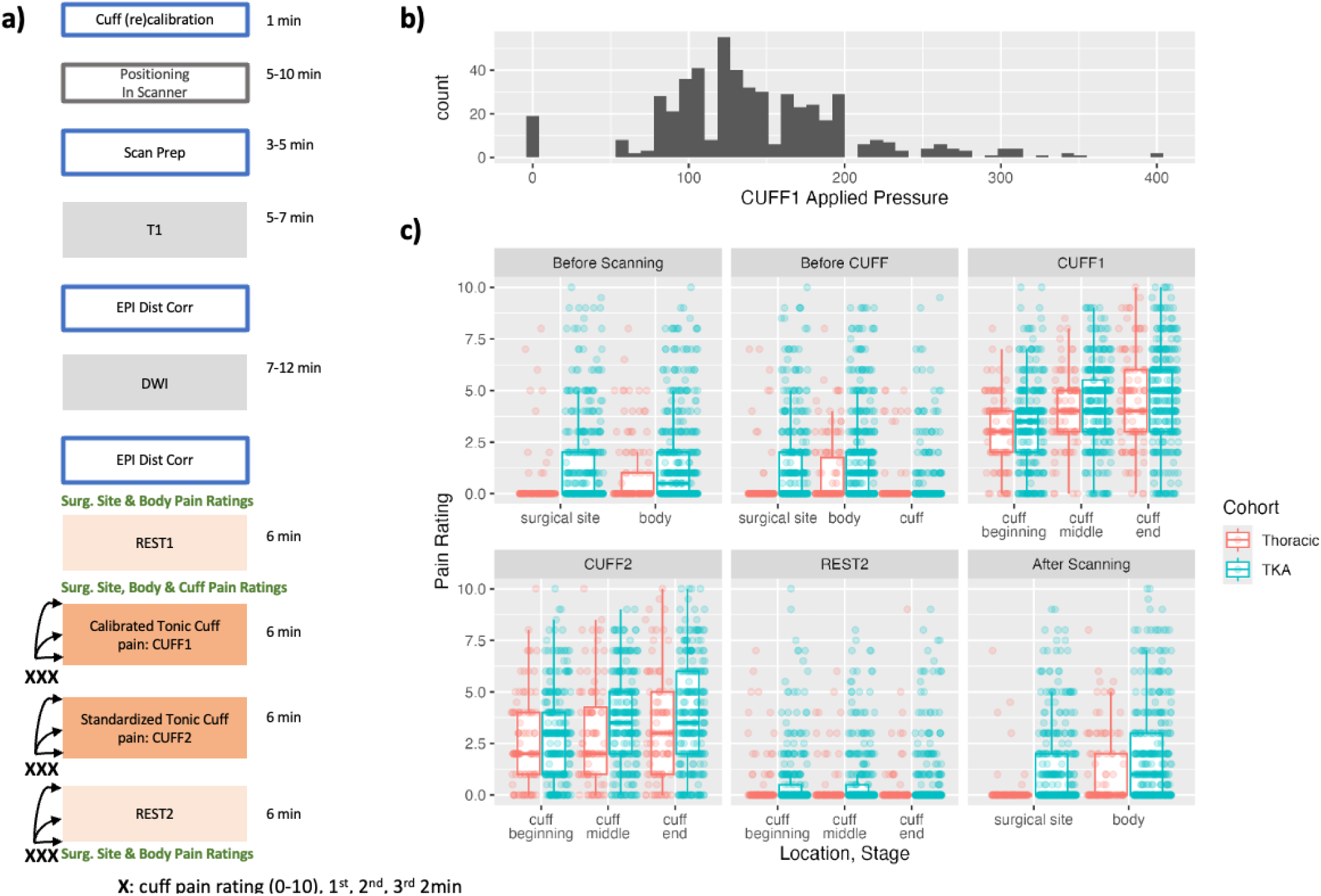
A2CPS Imaging Protocol. a) The protocol was designed for a one hour block of scanner time. The key portion consists of two fMRI scans acquired while the subject is wearing an inflated cuff, either set to a subject-specific (Calibrated) or fixed (Standard intensity) pressure; before and after the cuff scans, resting state fMRI is acquired, and T1-weighted, dMRI, and distortion correction images occur prior to the fMRI block. b) Distribution of Calibrated Pressures. c) Pain Ratings During tfMRI (CUFF) Scans.

The sMRI is a necessary input to all pipelines. However, several issues can degrade the quality of structural images (e.g., excessive motion). To mitigate the risk of low quality images, every sMRI is reviewed by the MRI technician during the session, who grades the image on a three-level scale (for details, see <u>Quality ratings</u>). The lowest rating is reserved for sMRIs that are expected to be unusable. Following this lowest rating, a second attempt is made to acquire an sMRI with a better rating. Only two attempts are made.

Prior to the fMRI scans, a pneumatic tourniquet cuff is placed on the subject’s leg. Inflating the cuff induces sustained pain (without damaging tissue), and the stimulus (pressure)-response function is conducive to both psychometric and neuroimaging study^40,41^. In the two resting state scans (REST1 and REST2), the cuff is uninflated. In the two tonic pain scans (CUFF1 and CUFF2), the cuff is inflated to constant pressures. The CUFF1 scans use a patient-calibrated constant pressure, which was determined as the pressure that elicits a pain rating of 4 on a 10 point scale. The CUFF2 scans used the same pressure across all participants (120 mm Hg). The minimal pressure for the individualized scans was 80 mm Hg, and if a lower pressure elicited pain higher than 4, the cuff scans and second rfMRI scan were excluded. In cases where inflation of the cuff is contraindicated, the first rfMRI scan was still acquired.

Acquisition harmonization was first carried out using test scans on phantoms. Then, prior to the start of the project, the same person was scanned at each MCC1 site. After MCC2 was added, another person was scanned at all MCC1 and MCC2 sites. All scans were covered by each local IRB using protocols for development scans.

In an effort to collect minimally processed raw data, no selectable scanner bias field or gradient distortion correction options are used (these options are not selectable on all scanners). All scans are collected without angulation (following the practice of the ABCD study^39^) because of concern about oblique scans on GE/Philips scanners.

Session data were collected and managed using Research Electronic Data Capture (REDCap)^42,43^. REDCap is a secure, web-based software platform designed to support data capture for research studies, providing 1) an interface for validated data capture; 2) audit trails for tracking data manipulation and export procedures; 3) automated export procedures for seamless data downloads to common statistical packages; and 4) procedures for data integration and interoperability with external sources.

### Acquired modalities

● **Anatomical (anat)**

○ **Structural MRI (sMRI)** provides information related to the anatomic layout of the brain, and allows calculations of cortical and subcortical volumes and morphology. It provides a stable reference for cross-subject and cross-modality alignments, and is thus an essential component.
● **Diffusion (dwi)**

○ **Diffusion-Weighted MRI (dMRI)** captures information about how the organization of brain tissues and structures hinders the natural diffusion of water molecules.
● **Functional (func)**

○ **Functional MRI (fMRI)** measures hemodynamically driven signal changes that result from localized changes in oxygen uptake and blood flow as a result of neural activity, and is typically analyzed by looking for signal variations associated with scan conditions (often controlled with a “task”) or at how spontaneous intrinsic variations in signal are coordinated across brain regions (“functional connectivity” of those areas).

▪ **Resting-state functional MRI (rfMRI)** are six minutes long and are done with a cuff, worn but uninflated.
▪ **Task-based functional MRI (tfMRI)** are also six minutes long and have a user calibrated inflation of the cuff (CUFF1) followed by a cuff inflation to a standard pressure (CUFF2).
● **Fieldmap (fmap)**

○ **Susceptibility-weighted MRI (swMRI)** enables susceptibility distortion correction. The A2CPS project acquires spin-echo B0 images with opposite phase encoding direction, which can be combined to create fieldmaps^44^. In general, the A2CPS B0 scans match the geometry of the target scans (i.e., the field of view and shimming of the scans to be corrected), though in some cases the geometry is slightly different.

### COVID-19

A challenging aspect of this project is that its data acquisition period (2021-present) overlapped with the COVID-19 pandemic. In addition to periods when MRI scanning was halted at individual sites based on local policies, institutional masking policies also varied across sites and across the period of the study, and in some cases required subjects to remain masked during all scans. After an initial adaptation period in mid-2021, individual mask usage in the scanner was recorded on a session by session basis. In that initial period, 170 sessions were conducted without specific recording of mask status.

## III. The A2CPS Processing Pipeline

To extract biomarkers from multisite neuroimaging data requires extensive curation, preprocessing, and modeling. Following the examples of many successful large projects that preceded A2CPS, including the Human Connectome Project^45^ and the UK Biobank^46^, we perform this extraction with automated processing pipelines built from software provided by the neuroimaging, data engineering, and A2CPS communities.

To reconstruct image data from *k*-space, each scanner’s manufacturer-specific algorithms and software are used (*k*-space data were not archived). The reconstructed images are exported in the Digital Imaging and Communications in Medicine (DICOM) format and sent electronically from participating sites to the Texas Advanced Computing Center (TACC). Transfers use encrypted communications channels and are stored with TACC’s Protected Data Service (corral-secure), which adheres to the Health Insurance Portability and Accountability Act.

At TACC, processing comprises three stages (Figure 3). First, the data are indexed and organized according to the Brain Imaging Data Structure (BIDS)^47^. Then, the data are sent through a series of pipelines, which were either drawn from the literature (as of the first release, this includes brainageR^48,49^, MRIQC^50^, fMRIPrep^51^, FreeSurfer^52^, QSIprep^53^, fsl_anat^54^, CAT12^55^, and NeuroMark^56^) or custom-built for A2CPS using several packages in the scientific computing literature (functional connectivity, calculated pain signature responses)^57–62^. Finally, the outputs from these pipelines are aggregated, de-identified, and stored alongside data from other modalities (e.g., demographics, genetic variant data, psychosocial measures).

**Figure 3.**
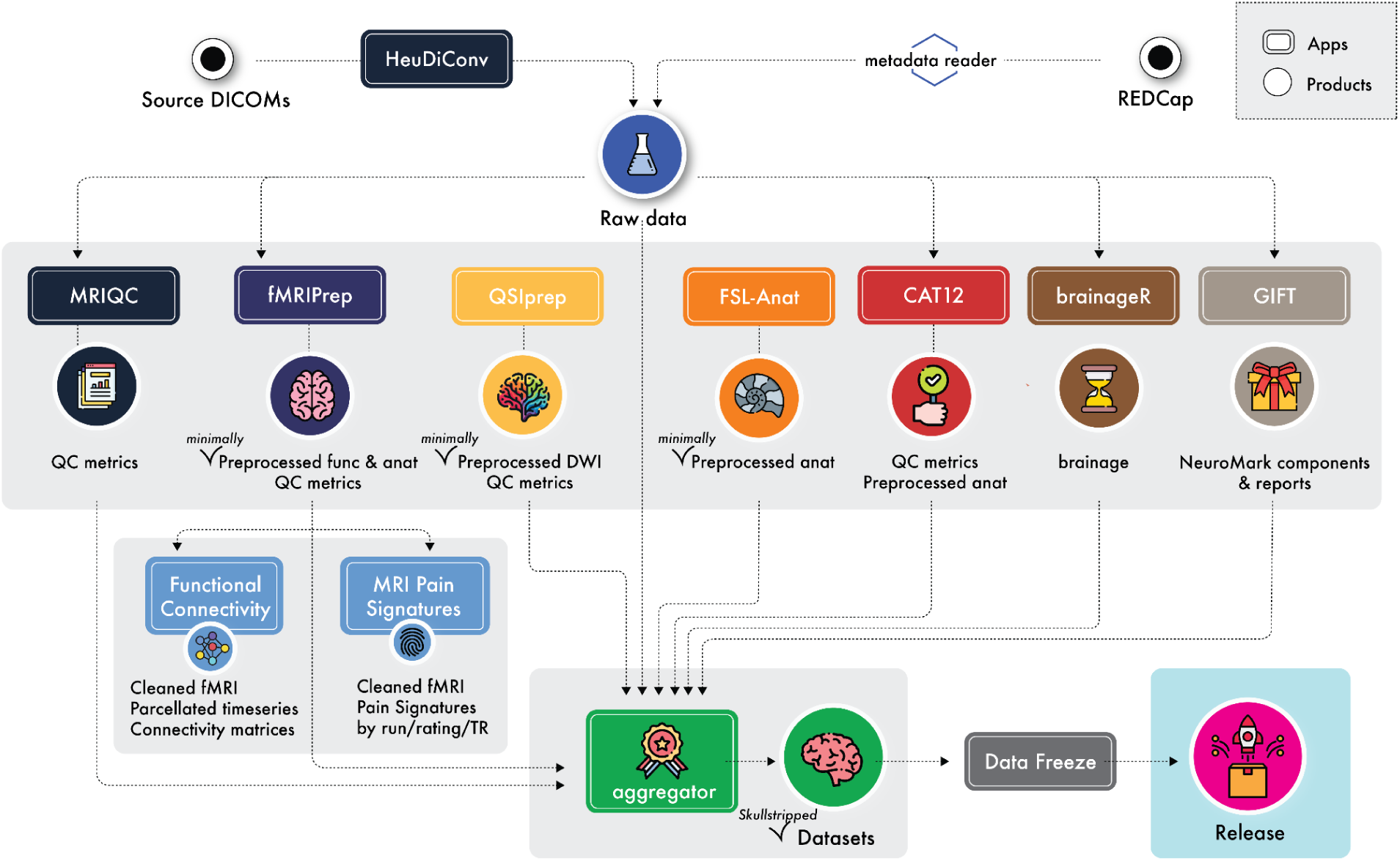
The A2CPS Image Processing Pipeline.

Processing is orchestrated with TACC Application Programming Interfaces (APIs; Tapis)^63^, which is a framework for developing web-based APIs to high-performance compute resources. Components of the A2CPS pipelines (e.g., subpipelines like MRIQC) are handled by Tapis “Apps”, and the Apps are coordinated by a set of Tapis Actors. Tapis Apps roughly correspond to BIDS Apps^64^ (that is, containerized^65,66^ code) which have been augmented with the capacity to run on TACC (e.g., wrapped in a Slurm^67^ script).

Following the Organization for Human Brain Mapping (OHBM) Committee on Best Practices in Data Analysis and Sharing (COBIDAS) report^68^, the DICOM files are converted into the Neuroimaging Informatics Technology Initiative (NIfTI) 1 format^69^ and organized according to the BIDS specification^47^. For this conversion, the pipeline uses HeuDiConv^70^, which in turn uses dcm2niix^71^. Conversion relies on the ReproIn^72^ heuristic file, modified to encode the sequence and file naming conventions developed for the A2CPS project. The original DICOMs are stored for posterity but are not included in data releases. The BIDS data are the most raw form of data (see also the section <u>Skull Stripping for Privacy</u>).

### Generation of Image Biomarkers

The first two aims of A2CPS require extraction of pre-specified measures that could serve as biomarkers or features of biosignatures. The generation of these measures guide preprocessing and analysis pipelines. Given that these measures were selected based on findings and proposals in the literature, the A2CPS pipeline replicates those published procedures, except in cases where procedures rely on pipelines that have been superseded.

The A2CPS Release 1.1.0 pipeline makes use of versions of six widely used processing streams (MRIQC 0.15.1^50,73^, fMRIPrep 20.2.3^51,74^, FreeSurfer 6.0.1^52^, `fsl_anat` 6.0.6.5^54,75^, CAT12 v12.8.1/r2042^55,76^, and QSIprep 0.21.4^53^, brainageR 2.1^48^, and GIFT 4.0.5.0/NeuroMark 2^56^). Note that many of these pipelines perform the same kinds of preprocessing steps (e.g., both MRIQC and fMRIPrep create brain masks), and in some cases do so with the same tools (e.g., both fMRIPrep and `fsl_anat` perform tissue-type segmentation with FSL’s `FAST` tool). No effort was made to combine steps across pipelines, and the products from all pipelines are archived, meaning that some products are duplicated. Duplicated products may not be exactly identical, as the preprocessing leading up to the step may diverge.

Pipeline products that are covered by BIDS adhere to that specification. Derivatives that have not yet been standardized by BIDS or an accepted BIDS extension proposal are stored in either a “BIDS-like” format (e.g., functional connectivity), or in the standard format produced by the pipeline (e.g., FreeSurfer, fsl_anat).

#### Preprocessing Overview

Minimally preprocessed derivatives for the anatomical, diffusion, and functional MRI data are produced by QSIprep and fMRIPrep. For detailed descriptions of the pipelines, see <u>Pipeline</u> <u>Boilerplate Text</u>. Anatomical scans underwent intensity inhomogeneity correction, skull-stripping, tissue segmentation, surface reconstruction, and spatial normalization. Diffusion images underwent denoising, Gibb’s unringing, head motion correction, eddy current correction, distortion correction, and normalization to standard space. Functional images underwent skull-stripping, susceptibility distortion correction, head motion correction, and slice-time correction. Both the diffusion and functional images were aligned to the anatomical image, and the anatomical and functional images are also provided in standard spaces.

#### Functional Connectivity

Functional connectivity profiles were estimated using custom pipelines. The inputs to this workflow were the preprocessed functional data from fMRIPrep. An overview of the pipeline is given in Figure 4. First, the initial 12 volumes (15 seconds) of each scan were trimmed, to allow for non-steady state normalization. Then, anatomical CompCor was run on the trimmed data, after bandpass filtering and *z*-scoring (Butterworth, 0.01 - 0.1 Hz)^1^. The top five components from the combined white matter and CSF mask along with the 24 motion parameters^77^ were used for nuisance regression (including the same trimming and filtering as when extracting the compcor components).

**Figure 4.**
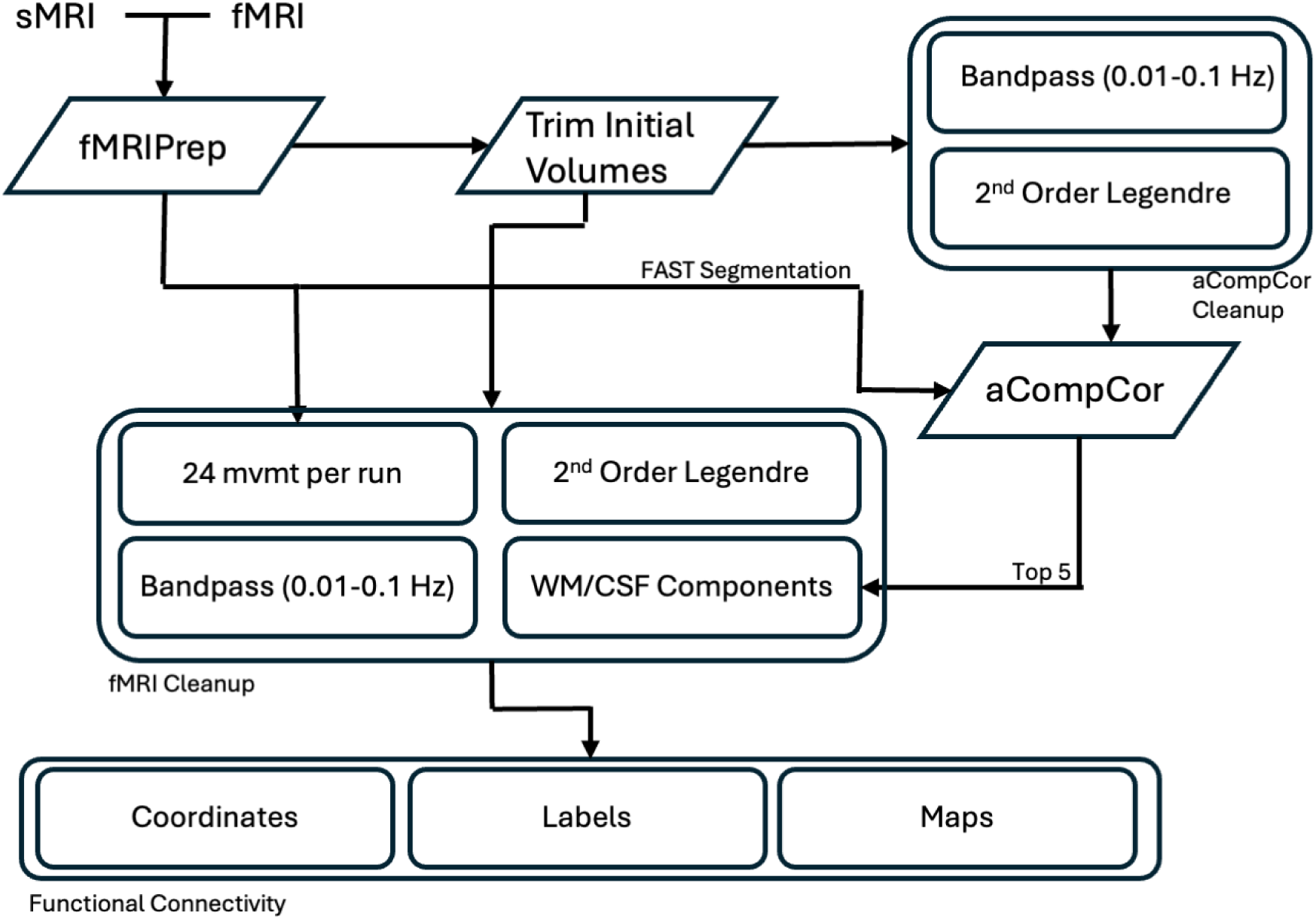
Functional Connectivity Workflow.

From the cleaned data, connectivity was estimated between three different kinds of regions: 1) nodes based on coordinates (10 mm sphere, Default Mode Network^20^, and a version of the Power atlas^78^, modified to exclude nodes within the cerebellum); 2) nodes based on probabilistic maps (64 and 1024 versions of the Dictionaries of Functional Modes^79^); and 3) standard, non-overlapping atlases (400 region Schaefer atlas^80^, Gordon atlas^81^, and a version of the Brainnetome^82^ atlas modified as in ^32^). Functional connectivity was estimated via the standard empirical covariance and the Ledoit-Wolf estimator^83^.

#### Neural Pattern Expressions

The pattern expressions from several multivariate models are thought to be related to the experience of pain and potentially predictive of chronic pain^17^, and so part of the A2CPS pipeline also involves extracting these pattern expressions. As with functional connectivity, this workflow received the preprocessed functional data from fMRIPrep, and the cleaning steps are similar (Figure 5). First, the initial 12 volumes (15 seconds) of each scan were trimmed, to allow for non-steady state normalization. First and second order polynomials were removed from the timeseries (that is, the mean was preserved), the 24 motion parameters^77^ were used for nuisance regression (including the same trimming and low-pass filtering at 0.1 Hz), and finally the cleaned data were Winsorized (trimming values exceeding 3 standard deviations from the mean).

**Figure 5.**
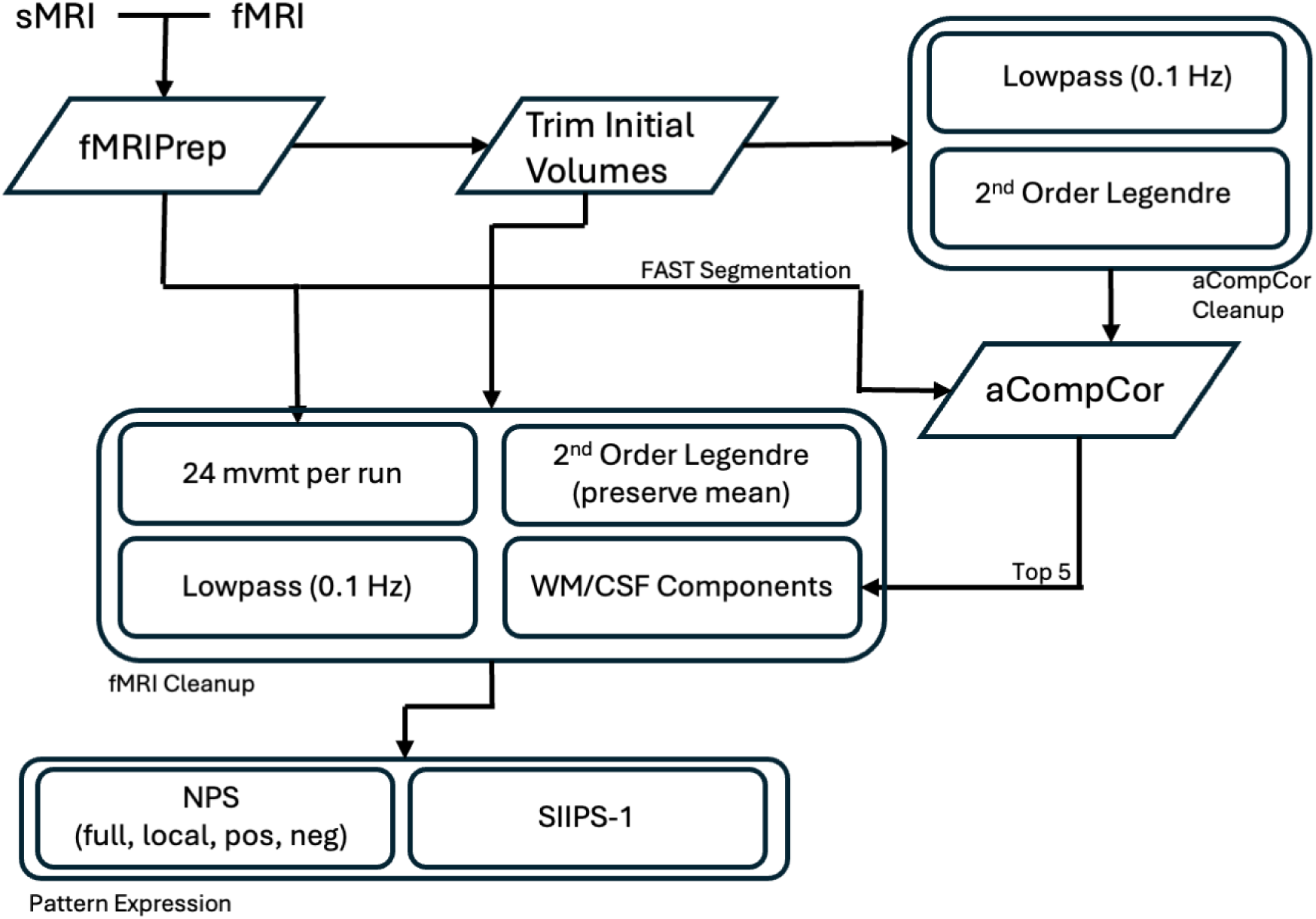
Neural Pattern Expression Workflow. NPS: Neurologic Pain Signature; SIIPS-1: Stimulus Intensity Independent Pain Signature 1.

After cleaning, pattern expressions were extracted. Extraction proceeded by calculating the similarity between a 3D volume and a standard mask. To enable flexibility in subsequent analyses, three measures of similarity were used: the dot product, cosine similarity, and product-moment correlation (for an overview of similarity measures considering one widely used pattern see ^84^). The volumes from which pattern expressions were extracted were also threefold: by individual TR, by the temporal average of two minute periods (corresponding to the three pain ratings provided by each participant), and by the runwise temporal average. For patterns, the following are provided: the neurologic pain signature (NPS)^85^, NPS regions (positive, negative, dorsal anterior cingulate cortex, left lateral occipital cortex, left superior temporal sulcus, posterior cingulate cortex, pregenual anterior cingulate cortex, right inferior parietal lobule, right lateral occipital cortex, right posterior lateral occipital cortex, left insula, right dorsal insula, right insula, right S2-Operculum, right thalamus, right V1, and vermis), and the stimulus intensity independent pain signatures-1 (SIIPS-1)^86^.

#### GIFT/NeuroMark

Intrinsic functional connectivity networks within the resting-state fMRI data were estimated with NeuroMark^56^ using the Group Independent Component Analysis (ICA) of fMRI Toolbox (GIFT; http://trendscenter.org/software/gift) BIDS app (version gift-bids:v4.0.5.4). NeuroMark is an automated and adaptive independent component analysis pipeline that uses group ICA-based templates as spatial priors to guide the identification of subject-specific independent component networks and the extraction of network features (e.g., timecourses, functional connectivity within and between networks, graph theoretic measures). We provide results from two different NeuroMark network templates. The first is NeuroMark 2.0 (model order = 175) which includes 58 non-artifactual components produced from a single model order of 175 components^87^. The second is the multi-scale NeuroMark 2.1 which includes 105 non-artifactual components covering diverse spatial scales which are aggregated from diverse model orders^87^.

Prior to NeuroMark, the resting-state fMRI data were minimally preprocessed (see Functional Data Preprocessing section), and then resampled to 2.4mm isotropic voxel size to ensure identical dimensions across all subjects. After resampling, volumes were spatially smoothed using a Gaussian kernel with a full width at half maximum (FWHM) of 6mm. NeuroMark was then applied separately for each resting-state run from each subject and separately for each template. The NeuroMark pipeline included the following configuration options: brain masking using the default mask plus intracranial volume, coregistration of the functional data to a functional (i.e., EPI) template in MNI space (separate from the registration performed during minimal preprocessing), removal of the first 15 volumes (i.e., dummy scans), bandpass filtering (0.01-0.15Hz), removal of the mean per timepoint, multivariate-objective optimization ICA with reference^88^, and *z*-score scaling of results. The NeuroMark outputs include, separately for each template applied to each resting-state run from each subject, the timecourses and spatial maps of each intrinsic component network, functional network connectivity estimates, summary reports, and visualizations of the networks.

### Skull Stripping for Privacy

Individual anatomic image outputs from all pipelines, as well as the source T1w images, are skull stripped prior to release (where not already skull stripped). The decision to apply this step was made in light of the growing concern that facial reconstruction from high resolution anatomical images might allow identification, conflicting with HIPAA’s requirement of removing “full-face photographs and any comparable images” (Privacy Rule, 45 CFR Part 164) to achieve de-identification. When brain masks were available (e.g., anatomical outputs of fMRIPrep), they were used to remove non-brain tissue. When brain masks were not available (primarily the raw data), skull-stripping was performed by application of a mask image generated by SynthStrip.^89^ All processing is carried out using unmodified raw images. Skull stripping removes useful information, making it impossible to extract some features ( e.g., intracranial volume). This loss is mitigated by provision of an array of features, which were calculated from the data before skull stripping (including intracranial volume).

Before opting for skull stripping, we considered more limited “defacing” approaches. There have been many efforts to target the face region specifically^90,91^, either by removing or altering signal. Defacing addresses some concerns about degrading image quality, making excising, blurring, or overlaying, rather than removing, the face region appealing. However, these less intrusive options leave a significant risk of identification^92,93^, since modern face recognition and reconstruction methods enable identification of images that have been defaced^91^. Moreover, altering even only the face region affects image processing relative to unchanged images^90,91,94^, and so analyses on images retain the skull but have been defaced will not necessarily produce the same results as analyses of raw images.

### Quality assurance and quality control

A major challenge with any large imaging project is assuring that acquisitions adhere to the expected setup, that no integrity problems occur (e.g. incomplete image exports or transfers), and that images are of sufficiently high quality. Our approach to quality relies on three tracks. The “fast” track performs an automated quality evaluation, and a “slow” one provides a more comprehensive evaluation that integrates manual review. A third, “*ad hoc*”, track includes checks that are not conducted with a set frequency but instead performed as needs arise (e.g., evaluation of acquisition changes necessitated by mandatory scanner software upgrades). All images that are flagged on Track 1 raw data checks are reviewed manually, but in general Track 2 evaluations do not occur on every scan. In addition to manually evaluating every scan flagged by Track 1 checks, we randomly sample from other scans as capacity permits.

#### Ongoing quality and integrity monitoring

The A2CPS DIRC monitors uploaded images and uses comparisons to patient records in REDCap to identify cases where imaging data might be missing or incomplete. The quality assurance process can potentially flag unexpected protocol deviations or errors in data collection.

Scans should result in BIDS data files with identical dimensions every time, and imaging parameters should be the same every time. These assumptions are tested using the BIDS validator^95^ as well as custom checks for variations in BIDS JSON values from each site’s reference scans. In addition to arising from use of an incorrect or modified protocol, unanticipated changes can potentially occur after a scanner upgrade, when settings specified as “default”, “shortest”, etc. in the saved configuration may lead to changed values.

Parameter checking is done on a subset of the fields in the JSON sidecars output during BIDS conversion, as well as dMRI .bval and .bvec files. Some numerical parameters are expected to exhibit small differences, so a magnitude of acceptable deviations for each parameter is defined. Scans that fail this check proceed through the rest of the analysis pipeline, but an automated warning is used to initiate manual review. Some failures have simple explanations (e.g., a functional scan may be short because it was terminated after the participant pressed the squeeze ball to request a stop), and others may involve extended communication with the imaging site.

#### Quality ratings

To facilitate monitoring the progress of the project and identifying problems in a timely manner, all A2CPS scans receive a quality rating on a three point “green”, “yellow”, “red” scale. Scans with a green or yellow rating are expected to be viable sources of biomarkers. Scans are marked yellow rather than green when they exhibit a minor quality issue that may warrant special consideration in some analyses. Scans that are marked red are not necessarily unusable, but biomarkers extracted from red scans are not expected to be comparable to typical ones.

The quality ratings are partially derived from image quality measures, which are obtained principally from MRIQC^50^,with others drawn from QSIPrep, fMRIPrep, and CAT12^55,96^.

##### Anatomical image

Labels are assigned initially as follows. By default, an image is marked as green. In Track 1, that rating can be downgraded in two ways:

1. The MRI technologist may label the image as red or yellow, following guidance and examples in an A2CPS technical manual.
2. An image may be downgraded from green to yellow if it receives a CAT12 Image Quality Rating below a relaxed threshold (80%), and further downgraded to red if the rating is below a stringent threshold (60%).
3. An image may be downgraded from green to red if CAT12 reports an Euler number of less than −217.^97^

The resulting rating for the image is therefore the lowest rating that was given by the technologist or CAT12. These ratings are then potentially refined in Track 2.

Track 2 manual reviews rely on the HTML reports produced by MRIQC, and those reports may be supplemented by the HTML report from fMRIPrep and the NiFTI image itself. The basic product of a manual review is a JSON file produced by the MRIQC “Rating Widget”. This widget provides a standardized list of artifacts and a summary of the image in a four category numerical rating (1: Exclude; 2: Poor; 3: Acceptable; 4: Excellent), which is collapsed into the three levels used by A2CPS (1: Red, 2,3: Yellow, 4: Green). This final summary rating is based on both (a) the worst rating from Track 1 (after confirming that the rating is not due to data entry error or correctable processing failure), and (b) the severity of the worst artifact present in the image.

##### Functional images

The MRI technologist does not rate these scans. Instead, functional images receive a default rating of “green” and are then potentially downgraded based on the following criteria:

1. Motion (following Parkes et al.^98^)

○ Images will be downgraded to yellow if the framewise displacement^99^ measures produced by MRIQC meet any of the following conditions

1. Average displacement > 0.25mm
2. More than 20% of the individual displacements are above a threshold (0.3mm for REST and 0.9mm for CUFF)
3. Any individual displacement value greater than 5mm
○ Images will be downgraded to red if the average framewise displacement is above 0.55mm
2. Integrity Checks

○ Truncated scans are always downgraded to red; this reflects the expectation that while they may be usable for many analyses, they are not strictly comparable to full length scans

All images receiving a rating of yellow or red due to motion are reviewed manually. These reviews are not expected to alter many ratings, but can confirm a lack of obvious errors in the Fast Track (e.g., inaccurate segmentation resulting in poor motion estimation). Additionally, a scan is manually reviewed if at least one MRIQC image quality measure is anomalous based on the distribution at that site.

##### Diffusion-Weighted Images

As with the other imaging modalities, the diffusion weighted images receive a default rating of “green”, and the rating is downgraded when integrity checks fail (e.g., when truncated scans were detected). Comparison of additional measures generated by QSIprep is in development.

#### QA phantom scans

Each site carries out regular QA phantom scans using a brief (<10 min) protocol, and these scans were used as a tool for coordinated monitoring of scanners by MCC physicists. The scans were also collated at TACC, which enables centralized processing and potentially useful extraction of long term trends that might inform analysis of study images^100^.

We run several measures, including the fBIRN QC regime^101^, on all such phantom scans. Though the main interest is in changes over time for each site, there is a high degree of standardization across sites – all (except UM, which uses the ABCD QA protocol) use a 17 cm diameter spherical volume (DSV) system phantom from GE or Siemens (UC, the sole Philips site, uses a GE phantom and other sites use one from the scanner manufacturer).

## IV. First A2CPS Data Release

The A2CPS project has opted for a phased release similar to that of ABCD and other projects. In the first release, scans and scan derivatives from pre-surgery visits completed through February 2023 will be available. The A2CPS project aimed to include only participants for which neuroimaging data could be collected, and known contraindications to receiving an MRI were an exclusion criteria for the study. However, several participants participated in the study but were not imaged due to a variety of reasons (i.e., unexpected contraindications, inability to schedule imaging prior to surgery). In total, the release includes 809 individuals of which 595 have imaging data (Table 3). Data for these participants is available through the National Institute of Mental Health Data Archive (NDA) (https://nda.nih.gov/).

**Table 3.** Demographics of A2CPS Participants with Imaging Data in Release 1.

| Characteristic | NS<br>N = 170<sup>1</sup> | SH<br>N = 23<sup>1</sup> | UC<br>N = 68<sup>1</sup> | UI<br>N = 179<sup>1</sup> | UM1<br>N = 75<sup>1</sup> | UM2<br>N = 60<sup>1</sup> | WS<br>N = 22<sup>1</sup> |
| --- | --- | --- | --- | --- | --- | --- | --- |
| Sex |  |  |  |  |  |  |  |
| Female | 99 (59%) | 15 (68%) | 50 (75%) | 108 (61%) | 44 (59%) | 34 (57%) | 10 (45%) |
| Male | 69 (41%) | 7 (32%) | 17 (25%) | 69 (39%) | 31 (41%) | 26 (43%) | 12 (55%) |
| Unknown | 2 | 1 | 1 | 2 | 0 | 0 | 0 |
| Age | 68 (63, 73) | 60 (56, 68) | 65 (59, 69) | 64 (58, 69) | 63 (54, 72) | 63 (56, 70) | 61 (52, 68) |
| Unknown | 3 | 3 | 2 | 2 | 0 | 1 | 3 |

<sup>1</sup>n (%); Median (Q1, Q3)

All 595 participants with imaging data have sMRI scans, but the remaining scans are more limited: dMRI: 589/595, REST1: 586/595, CUFF1: 458/595, CUFF2: 385/595, REST2: 477/595. The inflation of the cuff was contraindicated in some participants, resulting in termination of scanning after REST1, and in some additional cases, scan sessions were terminated early, usually at the request of the research participant. In rare cases, technical issues such as scan reconstruction failure on the scanner also reduced scan availability. There are three cases in which only the sMRI scan (the first in the session) is available.

Data derivatives only appear in this release if the entire A2CPS pipeline was completed without errors.

### Raw Data Quality

The quality ratings for images were designed such that only images with the lowest rating (“red”) will likely need to be excluded from most analyses. Considering each modality individually, approximately 98.7% of sMRI images, 99.8% of dMRI images, and 94.6% of fMRI images have a rating that is at least “yellow” or higher (Figure 6). For reference, note that the UK Biobank reported that, of the first 10,000 participants, 98.1% sMRI images passed their QC procedures, and only 81.1% of subjects passed QC on all scans (diffusion imaging was the worst, with 87.2% passing)^46^.

**Figure 6.**
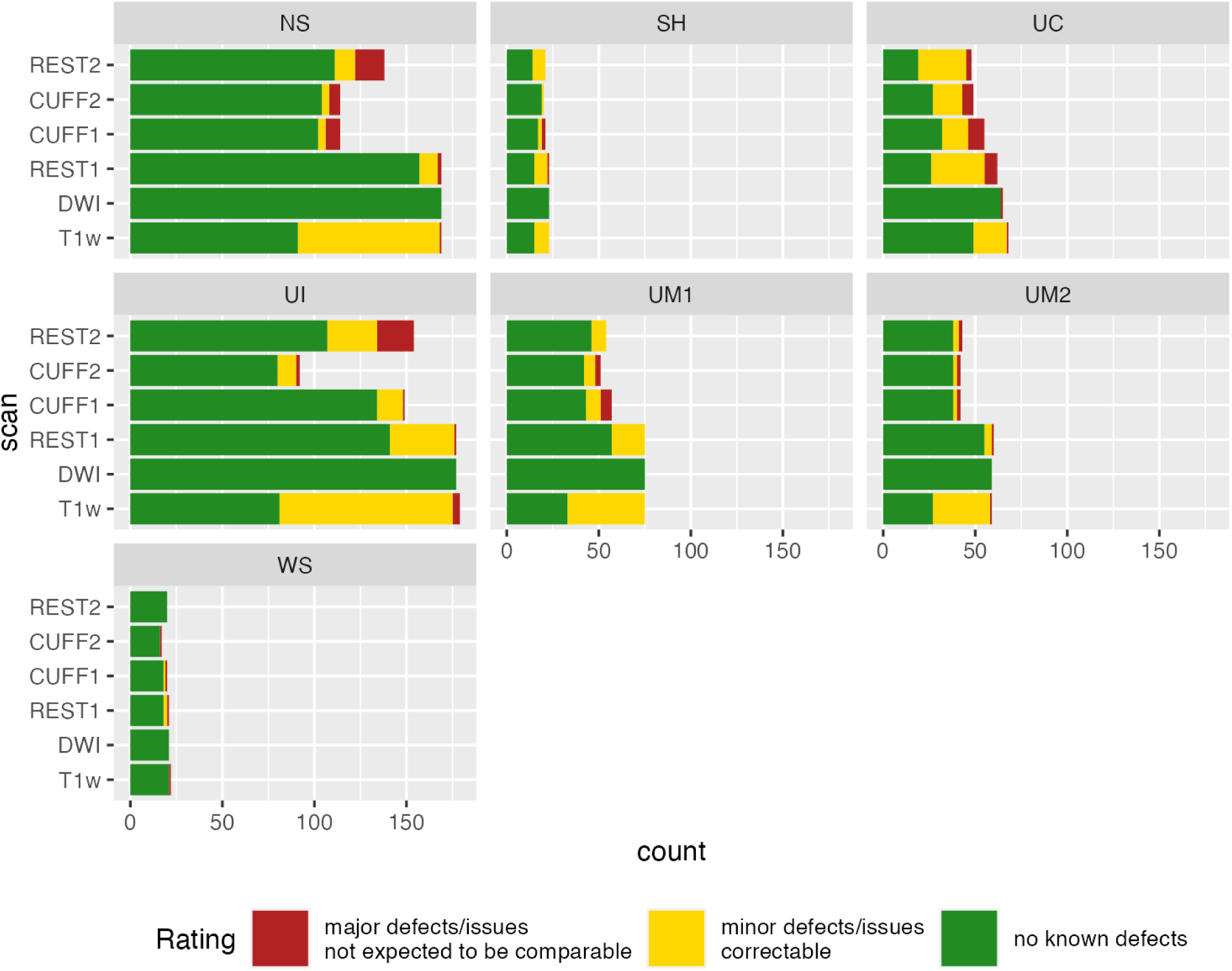
Distribution of Quality Ratings Across Scanners and Sites. All scans with a rating of “yellow” or “green” are expected to be usable for most analyses. Aggregated across sites, no scan type had more than 5% of scans with a rating of “red”.

One of the largest determinants of fMRI scan quality is in-scanner motion (see <u>Functional</u> <u>images</u>). In the first release, motion is comparable to that observed in the UKB, another large dataset that scanned a community (not MRI-trained) population (Figure 7a). On average, the difference in motion between the rfMRI and tfMRI scans was minimal, but motion in the tfMRI scans exhibited a longer tail (Figure 7b).

**Figure 7.**
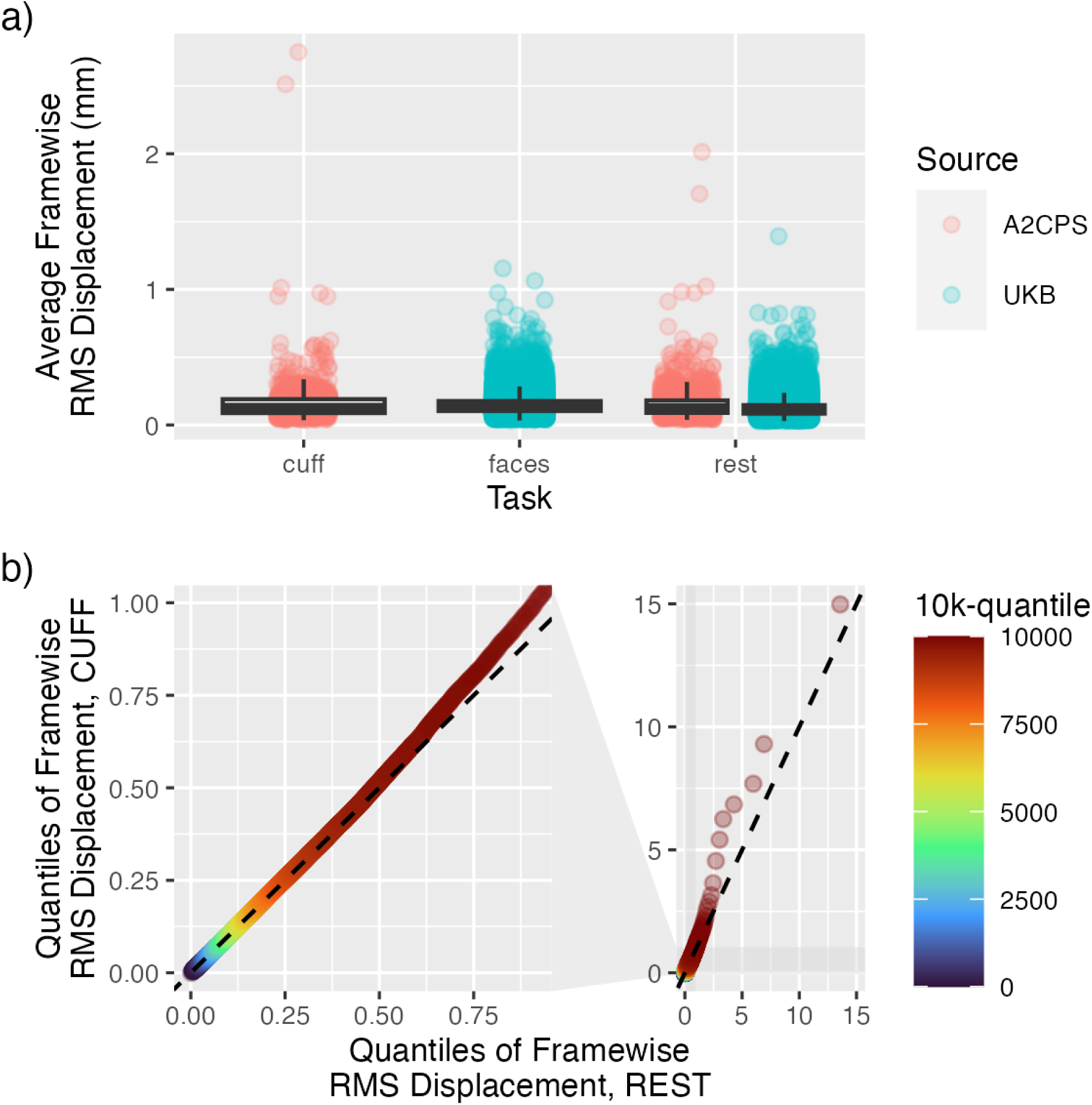
Motion in A2CPS Functional MRI. a) Overall Motion, Compared to External Dataset. The histograms show the distributions of average motion, normalized so that the largest bar has height of 1. Blue histograms are A2CPS, and the light gray histograms are the resting state scans of the UKB (same gray histogram in each panel). The point-intervals at the bottom show the median, 66% quantile interval, and 95% quantile interval of either the relevant spacetop task (blue) or UKB (black). b) Comparison of Framewise Motion Between REST and CUFF.

### Pressure and Pain Ratings

Early in data collection, CUFF1 scans were collected in which the applied pressure was below the standardized pressure (120 mmHg). After discussion within the consortium, it was decided that participants who provided a 4/10 pain rating on pressures below the one used during standardized scans (120 mmHg) would not receive any cuff pressure scans. Nevertheless, a subset of participants were collected before this rule was enacted, and these participants are included in the release (Figure 2b). Additionally, in a subset of scans, the cuff was not inflated. These cases are included in the main data release and the applied pressure is indicated as 0.

The cuff pressure effectively induced pain (Figure 2c). This is most pronounced in the thoracic cohort, which includes participants that tended to report no pain prior to the CUFF scans, pain during the CUFF scans, and no pain after the standardized pressure task. In general, there appears to be a trend whereby pain increases throughout CUFF scans.

### Image-Derived Phenotypes

In addition to the primary biomarkers, several derivatives are included in the release that may be of general interest. For example, as part of the A2CPS Pipeline, anatomical images are sent through FreeSurfer. All metrics provided by FreeSurfer (e.g., cortical thickness, mean curvature) are aggregated in tabular form. Given the size of the dataset, relationships between variables in the dataset may be detected with high power (e.g., with 600 participants there is 80% power to detect a correlation of 0.11 at the 0.05 significance level). For example, nine regional metrics provided by FreeSurfer show significant relationships with age (Figure 8).

**Figure 8.**
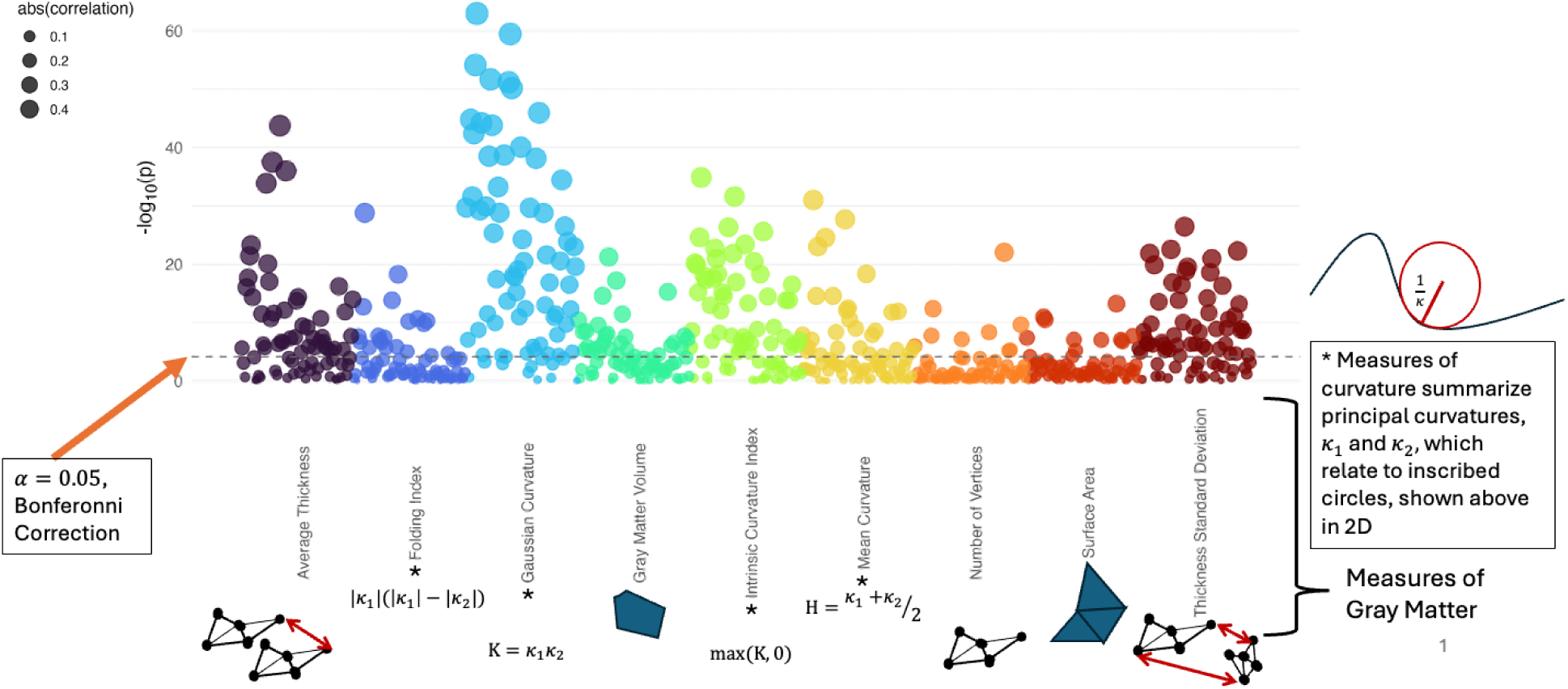
Relationships between Gray Matter Morphometry and Age. Points correspond to regions within the Destieux atlas^111^, with the height giving the statistical significance of a rank correlation between the morphometric measure and participant age. The dashed line marks significance at the 0.05 level after Bonferroni correction (across all measures and regions).

A key strength of the A2CPS dataset is its multimodality, with contents spanning psychosocial, quantitative sensory testing, electronic health records, genetic variants, ex-RNA, metabolomics, lipidomics, and proteomics as well as imaging. For example, one could use the data to study relationships between psychosocial variables and the brain age gap, which is a general marker of health that can be extracted from structural images (for review, see ^102^). Replicating previous findings, the brain age gap exhibits a relationship between depression^103,104^, anxiety^103^, and subjective cognitive function^105^ (Figure 9). In particular, a one point increase on the anxiety and depression scale corresponds to an increase in brain age by about 1.5 months, and a one point increase on the subjective cognitive function scale corresponds to a decrease by about 1.5 months (for detailed methods, see <u>Relationships Between Psychosocial Measures and Brain</u> <u>Age Gap</u>).

**Figure 9.**
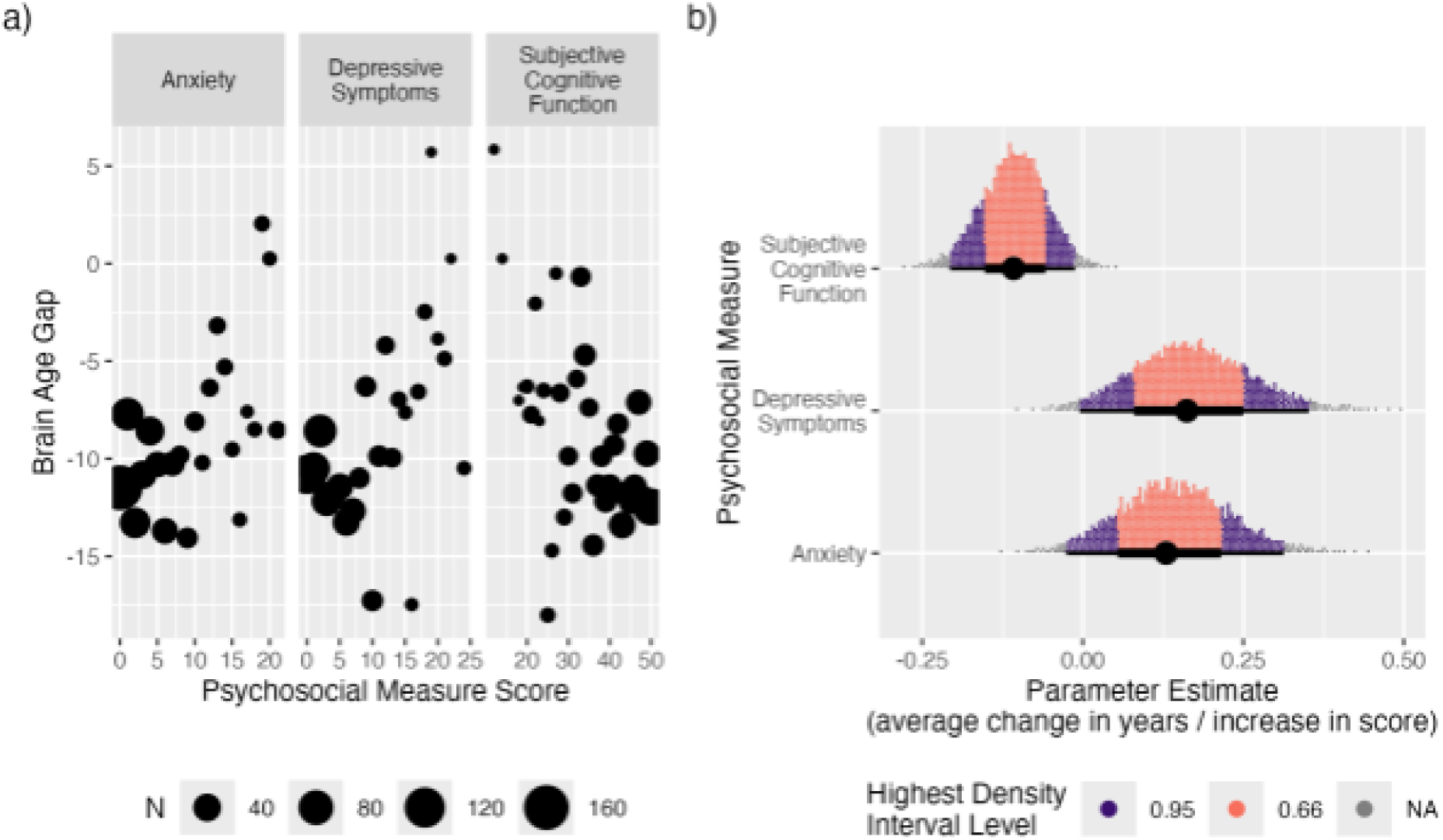
Relationships Between the Brain Age Gap and Psychosocial Measures. a) Average Gap at Measure Levels. At each score, the age gap was averaged. Point size corresponds to the number of participants in the corresponding averages. b) Highest Density Interval from Bayesian Parameter Estimation.

### Data Usage Agreement

Papers, book chapters, books, posters, oral presentations, and all other printed and digital presentations of results derived using the data should contain the following wording in the acknowledgments section:

> *Data were provided (in part) by the A2CPS Consortium funded by the National Institutes of Health Common Fund, which is managed by the OD/Office of Strategic Coordination (OSC). Consortium components include Clinical Coordinating Center (U24NS112873), Data Integration and Resource Center (U54DA049110), Omics Data Generation Centers (U54DA049116, U54DA049115, U54DA049113), MCC 1 (UM1NS112874), and MCC 2 (UM1NS118922).*

## V. Discussion and Future Work

The A2CPS research initiative provides a unique opportunity to study the transition to chronic pain in a large and diverse dataset. Here, we described the neuroimaging component of the A2CPS project – image acquisition, processing, biomarker extraction, and a summary of the currently available data. Neuroimaging provides a powerful source of objective markers that can contribute to improving patient outcomes. The dataset is expected to provide value both for pain researchers aiming to use image-derived phenotypes, and also for neuroimagers who wish to contribute to the ongoing challenge posed by chronic pain.

This paper focuses on the first of several planned releases. A2CPS has arranged a data release schedule that will rely on a form of semantic versioning (https://semver.org/), a set of three integers corresponding to major versions, minor versions, and version patches. Patches, if they prove necessary, will exclusively involve changes to documentation and bug fixes, but no additional content. Increments to minor versions may include additional derivatives or improvements to the processing pipeline (e.g., improved brain extraction algorithms). Major versions will additionally include new participants.

The first A2CPS data release includes the MRI data and products described here as well as other collected data, such as electronic health records and pain survey responses. Future releases will both add additional subjects and expand–or in some cases replace–the processing described in this report. In particular, subsequent releases may entail updating the processing pipelines. Any releases will be accompanied by documentation on updated processing.

Quality control during initial acquisition has focused on the raw data. We have focused on raw data to rapidly address any issues in protocols or procedures, and to provide collection sites feedback early in the data collection phase. As the project progresses, there will be additional quality control of the processing pipelines. Quality control of processing may entail both another quality rating scheme and also updates to the processing pipeline to incorporate improvements. For example, we have observed that FreeSurfer version 7 sometimes produces better outputs than the version 6 bundled with fMRIprep 20.2. In particular, in cases of a high degree of intensity non-uniformity, FreeSurfer 7’s use of the N4 algorithm^106^ instead of N3^107^ allows for better bias correction. These kinds of changes can occur in either minor or major release versions.

Although extensive work by the MCCs went towards harmonizing the scan protocols, which built on work by ABCD^39^, differences attributable to scanners are expected. The diversity of scanners, head coils, and multiband sequences used in the project can be viewed as both a weakness and a strength. The UK Biobank selected a single manufacturer (Siemens) and even a single scanner model (Skyra) for the brain imaging component of their project, as this eliminates sources of variability^108^. However, a diverse scanner population is not without benefits, as it facilitates characterization of measurements across a range of real-world imaging sources, facilitating generalizability. Diversity in image characteristics and other types of data has been crucial for enhancing performance across a range of artificial intelligence applications ^89,109,110^. We have observed varying levels of inter-scanner differences across different scans and measures, and the imaging products we have released should thus provide useful information when carrying out similar scans across different scanners. Future work will explore how to effectively combine data collected across scanners, accounting for confounding factors like differences in demographics at the scanning sites.

## Supplementary Materials

### Pipeline Boilerplate Text

Two of the key pipelines for generating minimally preprocessed images, QSIprep and fMRIPrep, provide boilerplate text describing their methods. The boilerplate text is copied, below.

#### QSIprep Processing

Preprocessing was performed using QSIPrep 0.21.4, which is based on *Nipype* 1.8.6^112,113^ (RRID:SCR_002502).

##### Anatomical data preprocessing

The T1-weighted (T1w) image was corrected for intensity non-uniformity (INU) using `N4BiasFieldCorrection`^106^ (ANTs 2.4.3), and used as T1w-reference throughout the workflow. The anatomical reference image was reoriented into AC-PC alignment via a 6-DOF transform extracted from a full Affine registration to the MNI152NLin2009cAsym template. A full nonlinear registration to the template from AC-PC space was estimated via symmetric nonlinear registration (SyN) using `antsRegistration`. Brain extraction was performed on the T1w image using `SynthStrip`^89^, and automated segmentation was performed using `SynthSeg`^114,115^ from FreeSurfer (7.3.1).

##### Diffusion data preprocessing

Any images with a b-value less than 100 s/mm^2 were treated as a b=0 image. Denoising using `patch2self`^116^was applied with settings based on developer recommendations. After `patch2self`, Gibbs unringing was performed using MRtrix3’s `mrdegibbs`^117^. Following unringing, the mean intensity of the DWI series was adjusted so all the mean intensity of the b=0 images matched across each separate DWI scanning sequence. B1 field inhomogeneity was corrected using `dwibiascorrect` from MRtrix3 with the N4 algorithm^106^ after corrected images were resampled.

FSL (version 6.0.5.1:57b01774)’s eddy was used for head motion correction and Eddy current correction^118^. Eddy was configured with a q-space smoothing factor of 10, a total of 5 iterations, and 1000 voxels used to estimate hyperparameters. A linear first level model and a linear second level model were used to characterize Eddy current-related spatial distortion. q-space coordinates were forcefully assigned to shells. Field offset was attempted to be separated from subject movement. Shells were aligned post-eddy. Eddy’s outlier replacement was run^119^. Data were grouped by slice, only including values from slices determined to contain at least 250 intracerebral voxels. Groups deviating by more than 4 standard deviations from the prediction had their data replaced with imputed values.

Data was collected with reversed phase-encoding blips, resulting in pairs of images with distortions going in opposite directions. Here, b=0 reference images with reversed phase encoding directions were used along with an equal number of b=0 images extracted from the DWI scans. From these pairs the susceptibility-induced off-resonance field was estimated using a method similar to that of `topup`^120^. The fieldmaps were ultimately incorporated into the Eddy current and head motion correction interpolation. Final interpolation was performed using the `jac` method.

Several confounding time-series were calculated based on the preprocessed DWI. These included framewise displacement^121^, which was calculated using the implementation in *Nipype*. The head-motion estimates calculated in the correction step were also placed within the corresponding confounds file. Slicewise cross correlation was also calculated.

The DWI time-series were resampled for alignment with the anterior commissure - posterior commissure line, generating a preprocessed DWI run in ACPC space with 1.7mm isotropic voxels.

Many internal operations of *QSIPrep* use *Nilearn* 0.10.1^122^ ( RRID:SCR_001362) and *Dipy*^123^.

For more details of the pipeline, see the section corresponding to workflows in *QSIPrep*’s documentation: https://qsiprep.readthedocs.io/en/latest/workflows.html.

#### fMRIPrep Processing

Anatomical, resting, and task MRI preprocessing was performed using fMRIPrep, which is based on Nipype 1.6.1^112,113^ (RRID:SCR_002502).

##### Anatomical data preprocessing

The T1-weighted (T1w) image was corrected for intensity non-uniformity (INU) with `N4BiasFieldCorrection`^106^, distributed with ANTs 2.3.3^124^ (RRID:SCR_004757), and used as T1w-reference throughout the workflow. The T1w-reference was then skull-stripped with a Nipype implementation of the `antsBrainExtraction.sh` workflow (from ANTs), using OASIS30ANTs as target template. Brain tissue segmentation of cerebrospinal fluid (CSF), white matter (WM) and gray matter (GM) was performed on the brain-extracted T1w using `fast`^125^ (FSL 5.0.9, RRID:SCR_002823). Brain surfaces were reconstructed using `recon-all`^126^ (FreeSurfer 6.0.1, RRID:SCR_001847), and the brain mask estimated previously was refined with a custom variation of the method to reconcile ANTs-derived and FreeSurfer-derived segmentations of the cortical gray matter of Mindboggle^127^ (RRID:SCR_002438). Volume-based spatial normalization to two standard spaces (MNI152NLin2009cAsym, MNI152NLin6Asym) was performed through nonlinear registration with `antsRegistration` (ANTs 2.3.3), using brain-extracted versions of both T1w reference and the T1w template. The following templates were selected for spatial normalization: *ICBM 152 Nonlinear Asymmetrical template version 2009c*^128^ (RRID:SCR_008796; TemplateFlow ID: MNI152NLin2009cAsym), FSL’s *MNI ICBM 152 non-linear 6th Generation Asymmetric Average Brain Stereotaxic Registration Model*^129^ (RRID:SCR_002823; TemplateFlow ID: MNI152NLin6Asym).

##### Functional data preprocessing

For each of the BOLD runs found per subject, the following preprocessing was performed. First, a reference volume and its skull-stripped version were generated using a custom methodology of fMRIPrep. A B0-nonuniformity map (or *fieldmap*) was estimated based on two (or more) echo-planar imaging (EPI) references with opposing phase-encoding directions, with `3dQwarp`^130^ (AFNI 20160207). Based on the estimated susceptibility distortion, a corrected EPI (echo-planar imaging) reference was calculated for a more accurate co-registration with the anatomical reference. The BOLD reference was then co-registered to the T1w reference using `bbregister` (FreeSurfer) which implements boundary-based registration^131^. Co-registration was configured with nine degrees of freedom to account for distortions remaining in the BOLD reference. Head-motion parameters with respect to the BOLD reference (transformation matrices, and six corresponding rotation and translation parameters) are estimated before any spatiotemporal filtering using `mcflirt`^99^ (FSL 5.0.9). BOLD runs were slice-time corrected using `3dTshift` from AFNI 20160207^130^ (RRID:SCR_005927). The BOLD time-series were resampled onto the following surfaces (FreeSurfer reconstruction nomenclature): *fsaverage*. The BOLD time-series (including slice-timing correction when applied) were resampled onto their original, native space by applying a single, composite transform to correct for head-motion and susceptibility distortions. These resampled BOLD time-series will be referred to as preprocessed BOLD in original space, or just preprocessed BOLD. The BOLD time-series were resampled into standard space, generating a preprocessed BOLD run in *MNI152NLin2009cAsym* space. First, a reference volume and its skull-stripped version were generated using a custom methodology of fMRIPrep. Grayordinates files^45^ containing 91k samples were also generated using the highest-resolution *fsaverage* as intermediate standardized surface space. Several confounding time-series were calculated based on the preprocessed BOLD: framewise displacement (FD), DVARS and three region-wise global signals. FD was computed using two formulations following Power (absolute sum of relative motions^121^) and Jenkinson (relative root mean square displacement between affines^99^). FD and DVARS are calculated for each functional run, both using their implementations in Nipype (following the definitions by [56]). The three global signals are extracted within the CSF, the WM, and the whole-brain masks. Additionally, a set of physiological regressors were extracted to allow for component-based noise correction (*CompCor*^132^). Principal components are estimated after high-pass filtering the preprocessed BOLD time-series (using a discrete cosine filter with 128s cut-off) for the two *CompCor* variants: temporal (tCompCor) and anatomical (aCompCor). tCompCor components are then calculated from the top 2% variable voxels within the brain mask. For aCompCor, three probabilistic masks (CSF, WM and combined CSF+WM) are generated in anatomical space. The implementation differs from that of Behzadi et al. in that instead of eroding the masks by 2 pixels on BOLD space, the aCompCor masks are subtracted from a mask of pixels that likely contain a volume fraction of GM. This mask is obtained by dilating a GM mask extracted from the FreeSurfer’s *aseg* segmentation, and it ensures components are not extracted from voxels containing a minimal fraction of GM. Finally, these masks are resampled into BOLD space and binarized by thresholding at 0.99 (as in the original implementation). Components are also calculated separately within the WM and CSF masks. For each CompCor decomposition, the *k* components with the largest singular values are retained, such that the retained components’ time series are sufficient to explain 50 percent of variance across the nuisance mask (CSF, WM, combined, or temporal). The remaining components are dropped from consideration. The head-motion estimates calculated in the correction step were also placed within the corresponding confounds file. The confound time series derived from head motion estimates and global signals were expanded with the inclusion of temporal derivatives and quadratic terms for each^133^. For all functional scans, frames that exceeded a threshold of 1.5 standardized DVARS were annotated as motion outliers. For CUFF scans, frames that exceeded 0.9 mm FD were also annotated, and for REST the FD threshold was 0.3 mm. All resamplings can be performed with a single interpolation step by composing all the pertinent transformations (i.e. head-motion transform matrices, susceptibility distortion correction when available, and co-registrations to anatomical and output spaces). Gridded (volumetric) resamplings were performed using `antsApplyTransforms` (ANTs), configured with Lanczos interpolation to minimize the smoothing effects of other kernels^134^. Non-gridded (surface) resamplings were performed using `mri_vol2surf` (FreeSurfer).

Many internal operations of fMRIPrep use Nilearn 0.6.2^122^ (RRID:SCR_001362), mostly within the functional processing workflow. For more details of the pipeline, see <u>the section</u> <u>corresponding to workflows in fMRIPrep’s documentation</u>.

### Active Elements

Three of the Siemens sites employ a 64-Channel Head Coil (Table 2; NS, WS, RU). Although up to 64 channels can be used, the scanners have automated systems for determining which channels should be active based on a participant’s positioning and the placement of the imaging volume relative to the coil elements. These automated systems were used, meaning that the selection of active coil elements was variable across participants; additionally, given that scan technicians sometimes needed to reposition the scan volume because participants could adjust themselves between scans, in rare cases coil elements used can even vary across scans within a single session. This variable selection of active elements could influence the signal-to-noise ratio across participants, particularly in the cortex. In practice, the variability in signal-to-noise within gray matter, across participants, was not substantially lower than the equivalent metric from images collected by other scanners (Figure S1). Where relevant, the active elements are recorded in the raw bids dataset.

**Figure S1.**
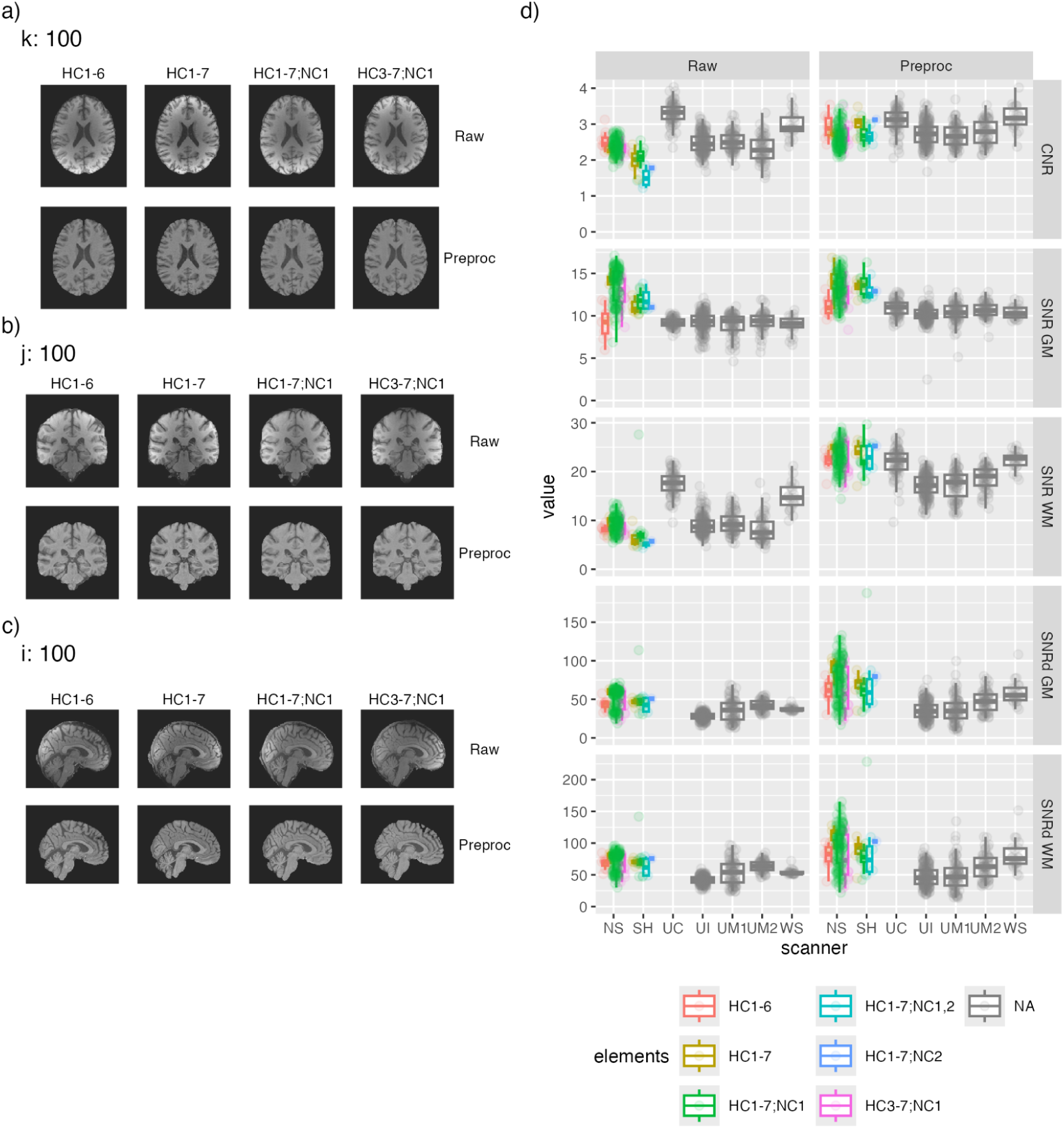
Variable Active Element Selection. a-c) Representative Participants from NorthShore with Varied Element Selection, Before (Raw) and After (Preproc) Preprocessing. d) Quality Metrics, Before (Raw) and After (Preproc) Preprocessing. Note that the SNRd values from the scanner at UC are omitted (this scanner results in most background voxels with intensity zero, which results in nonsensical values for this method of calculating signal-to-noise).

### Use of SENSitivity Encoding with Philips

Most of the anatomical images were collected with partially parallel acquisitions that fill missing data in k-space (e.g.,generalized autocalibrating partially parallel acquisitions; GRAPPA^135^), whereas the images collected on the Philips scanner at the University of Chicago were collected with a technique that unmixes and combines in image space (SENSitivity Encoding; SENSE^136^). Images reconstructed with SENSE have a background intensity that is often near zero (Figure S2). The zero-value background voxels precludes some analyses and heuristics, such as the estimation of image noise from regions of minimal signal.^137^

**Figure S2.**
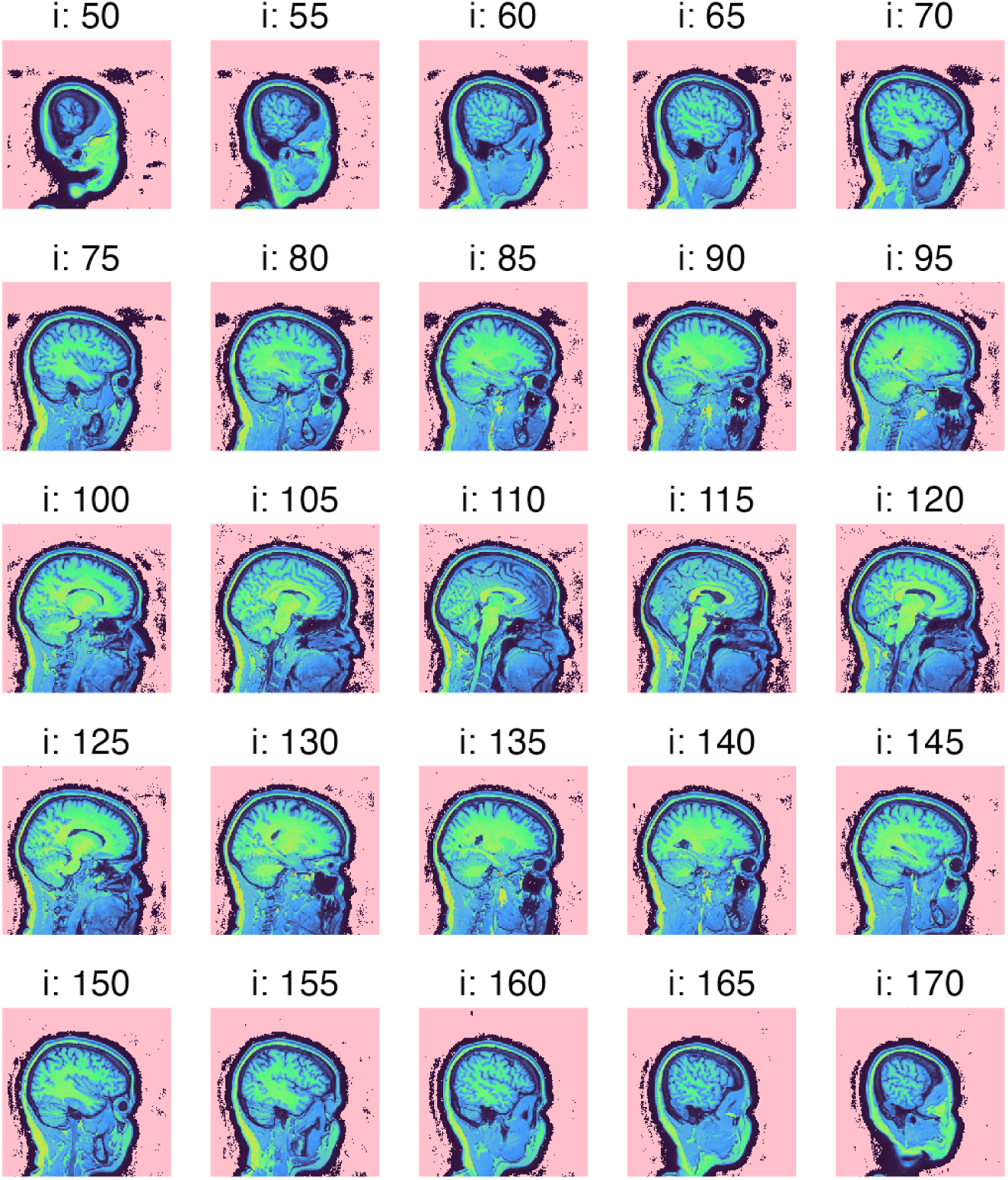
Representative Anatomical Image Collected with SENSE. For display, voxels with exactly zero intensity are colored in pink. Most voxels in the background are exactly zero.

### Impact of Limited Field-of-View at Sites

The ABCD acquisition protocol specifies that the functional scans should be collected with 60 slices, but that size was not feasible at two of the MCC1 sites: NS and UC. Instead, those sites reduced the slice stack to 54. This restriction meant that the coverage differed across sites. In particular, functional scans from NS and UC excluded the cerebellum at a higher rate than other sites (Figure S3); in NS scans, approximately 25% of the cerebellum was excluded, and in UC scans, approximately 12% of the cerebellum was excluded.

Although the limited field-of-view primarily affected acquisition of the cerebellum, individual scans also excluded portions of the cerebrum.

**Figure S3.**
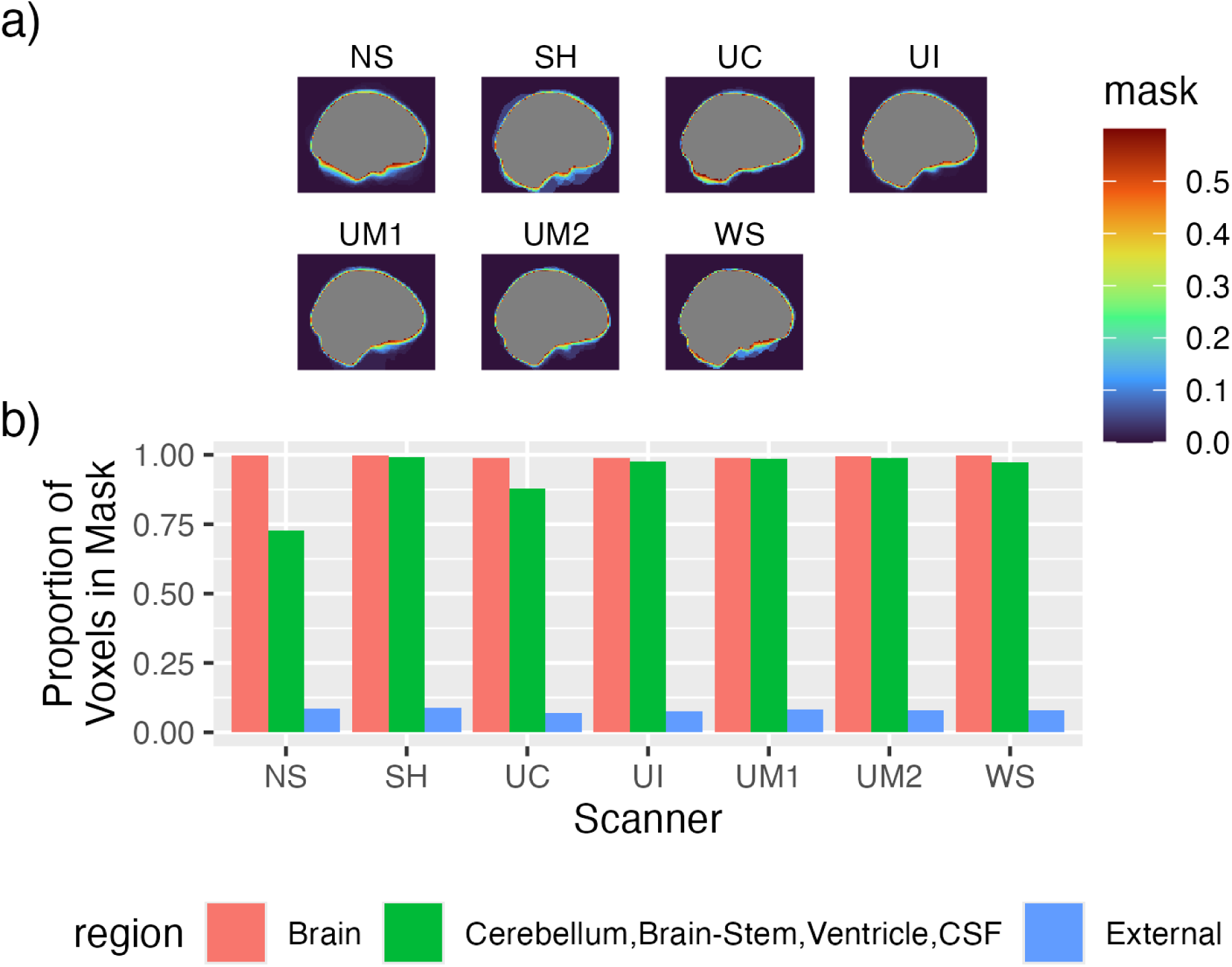
Field-of-View Across Scanners. a) Proportion of Voxels in Mask, by Scanner. Proportions calculated across REST1 scans (in MNI152NLin2009cAsym space). Note that the scale is clipped at 0.6. b) Proportion of Voxels in Mask by Region and Site. Heights correspond to the proportion of voxels that are in the intersection of a brain mask and a given region within a REST1 scan, out of all voxels that are in that region for the same kind of scan.

### Non-Steady State Scans

In most fMRI scans, an initial set of non-steady state volumes were reported by MRIQC, which bases that detection on the Nipype interface `NonSteadyStateDetector`. That interface marks as non-steady state the first contiguous sequence of volumes that are outliers – volumes whose average signal has a modified z-score larger than 3.5 ^138^ (based on only the initial 50 volumes). This criterion selects a variable number of volumes across functional runs (Figure S4). For ease of downstream analyses, a fixed count of 15 volumes (equivalently, 12 seconds) was applied.

**Figure S4.**
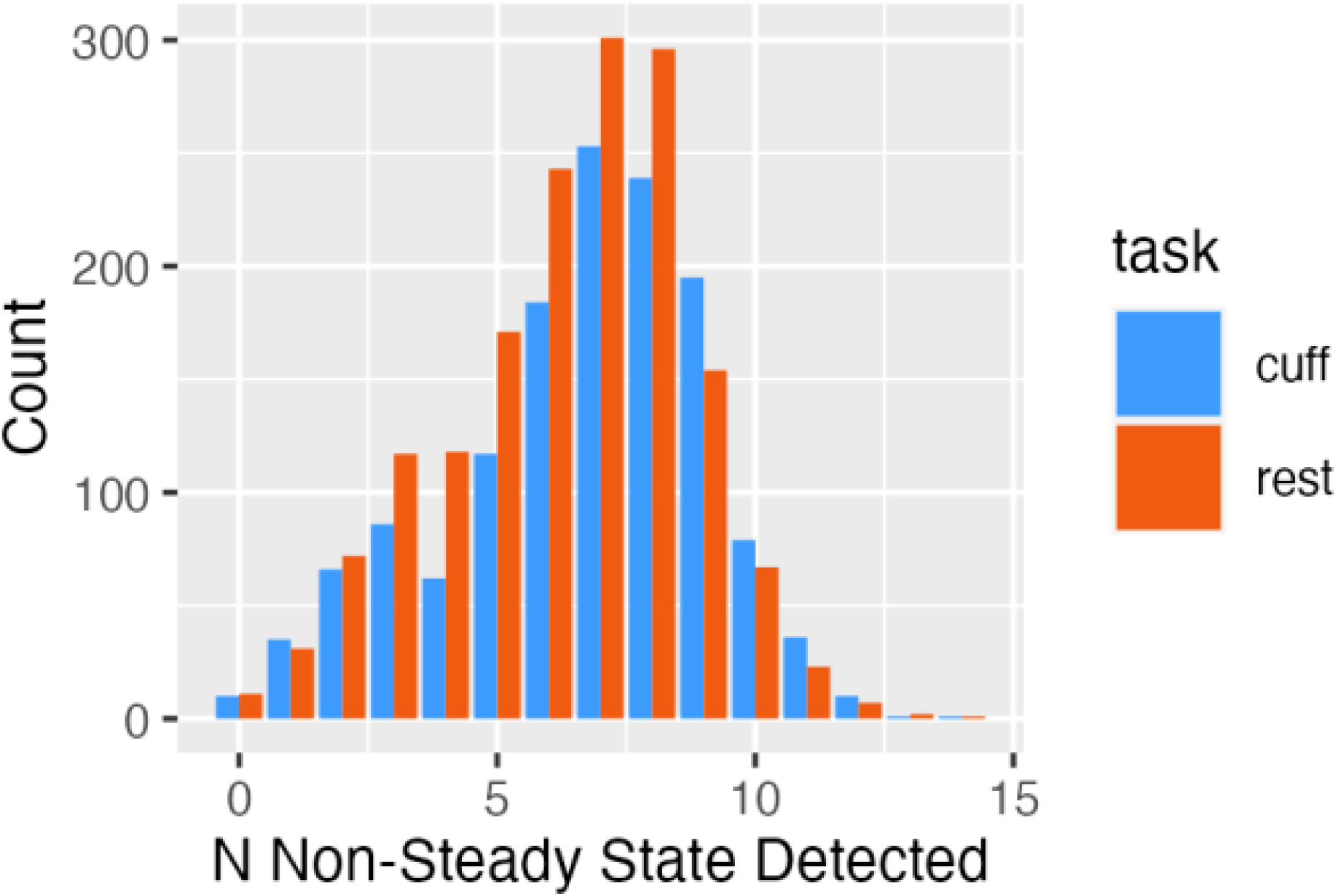
Number of Non-Steady States in Functional Scans. Counts correspond to scans.

### Standard Pipeline Modifications

Some of the pipelines from the literature were unable to process A2CPS participants using default settings. Pipeline failures were monitored, and in some cases modifications allowed for successful processing. These modifications are documented here.

#### fsl_anat

##### Field-of-View Estimation in UC

An early step in this pipeline involves estimating the field-of-view and cropping the image to a standard size. The cropping is done with the FSL tool `robustfov`, which uses a heuristic to first identify the most superior slice that could include the skull, uses a pre-specified distance in mm to select an inferior slice that is expected to be below the cerebellum, and crops everything outside of those slices. The heuristic is based on the quantiles of intensity values, but this heuristic was observed to often fail for data collected with SENSE.

To accommodate scans that failed for this reason, a small amount of noise was added to the entire image, and then the field-of-view was estimated on this degraded image. The field-of-view was then applied to the original image, and that cropped image was provided to fsl_anat (with automated cropping disabled).

##### High-Intensity Voxels

A subset of cases exhibited poor alignment to the standard space, with the misalignment driven by high-intensity voxels from outside of the brain. The extreme intensity values may have been driven by relatively high proportions of fatty tissue. For cases where these failures were identified, a mask was generated from the top 1% of voxels, which was then visually inspected to confirm that it did not include any voxels in the brain. That mask was then provided to fsl_anat as a “lesion” mask (via a hidden option), resulting in those voxels being ignored during alignment. In instances where the mask included brain voxels, the `fsl_anat` results were not included.

#### brainageR

##### Oblique Datasets

The brainageR pipeline would not process images that were oblique. Oblique datasets were detected with `nibabel` and reoriented to cardinal axes with AFNI’s `3dWarp`.

### Relationships Between Psychosocial Measures and Brain Age Gap

As part of the first data release, we explored relationships between psychosocial measures and the brain age gap. The gap was calculated using `brainageR`^49^, deployed in a custom docker image. The age gap was related to three psychosocial measures calculated in the first release: Depression (Patient Health Questionnaire Depression Scale^139^), Anxiety (Generalized Anxiety Disorder 7 Item Scale^140^), and subjective cognitive function (Multidimensional Inventory of Subjective Cognitive Impairment^141^). First, for visualization (Figure 10a), the gap was averaged at each level of the scale. Second, the scales were used as continuous predictors in a Bayesian regression model (including regressors for age, sex, an interaction between age and sex, and scanner), which was fit using `brms`^142^. To estimate the posterior, 8 chains were run for 2000 iterations each with the first 1000 reserved for warmup. The priors were left as defaults, excepting on the intercept and regression coefficients for which N(0,10) were used. All posteriors had tail effective sample size over 4000 and 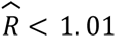.^143^

## Footnotes

1 Although fMRIPrep provides aCompCor components, those components are extracted from data that have been temporally filtered in a way that is not configurable.

